# Infarctsize-AI: an efficient infarct size image analysis tool for small rodent myocardial infarction studies

**DOI:** 10.1101/2025.11.18.688527

**Authors:** Csenger Kovácsházi, Dóra Kapui, Bennet Y Weber, Tamás G Gergely, Gábor B Brenner, Bence Ágg, Csanád Tabajdi, Adrienn Rácz, András Horváth, Sauri Hernandez-Resendiz, Derek J Hausenloy, Reinis Vilskersts, Marta Oknińska, Michał Waszkiewicz, Michal Mączewski, Arnold Molnár, Tamara Szabados, Péter Bencsik, Thomas Krieg, Javier Inserte, Rainer Schulz, Coert J Zuurbier, Ioanna Andreadou, Bruno K Podesser, Péter Ferdinandy, Zoltán Giricz

## Abstract

**Background:** Myocardial infarct size (IS) is the gold standard end-point in shorth-term studies on cardioprotection. However, IS quantification in rodent models with standard Evans Blue and 2,3,5-triphenyltetrazolium chloride (TTC) staining is time-consuming and prone to inter-observer variance. Therefore, we aimed to develop an artificial intelligence (AI)-based application to reduce time and inter-observer variability of IS analysis in rodent acute myocardial infarction (MI) models.

**Methods:** We used TTC/Evans blue-stained heart slice images of independent laboratories from previously published projects. Rat (n = 325 and 248 slices) and mouse (n = 77 slices) datasets were used to train deep learning segmentation models with three different neural network architectures, which were combined into a single AI analysis.

AI analysis was compared with manual analysis on rat data from a training laboratory (internal data, n = 496 slices, n = 41 whole-hearts) and data from independent laboratories (external data, n = 60 and 62 slices). Additionally, two independent evaluators performed manual and AI-assisted analysis, consisting of AI-analysis and its manual correction, on internal (n = 36 slices) and external data (n = 37 slices).

**Results:** Lin’s concordance correlation coefficient (CCC) between IS/AAR values from manual and AI analysis was 0.844 with 95% CI of [0.814; 0.869] for images of internal data heart slices. On external data heart slices, AI accurately annotated slice area and AAR but failed to annotate infarcted area. On internal whole-heart data, CCC between AI and AI-assisted IS/AAR was 0.894 with 95% CI of [0.812; 0.942]. AI-assisted analysis reduced evaluation time on both internal and external datasets and increased region overlap for AAR between the two independent evaluators on dependent data.

**Conclusions:** AI-assisted analysis significantly reduced analysis time and inter-observer variability. For optimal performance, lab-specific AI training is recommended. Infarctsize-AI™ is available at https://infarctsize.com.

**Translational perspective:** Myocardial infarct size (IS) is the gold-standard end-point in shorth-term studies to assess potential cardioprotective therapies against acute myocardial infarction (AMI). However, IS quantification in rodent AMI models is time-consuming and prone to inter-observer variance. Therefore, we developed an AI-based software that can reduce analysis time and inter-observer variability and facilitate documentation, which facilitates the clinical translation of potential cardioprotective therapies.

**Graphical abstract:** 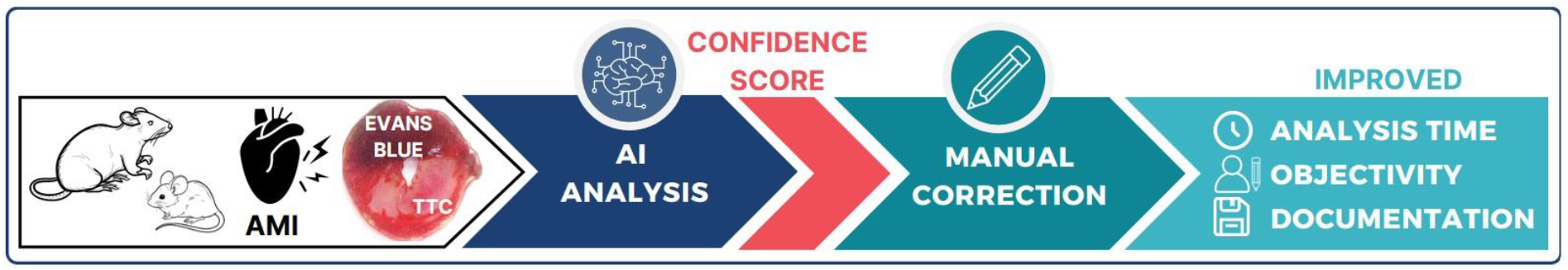

## Introduction

Myocardial infarct size (IS) continues to be a leading predictor of adverse cardiac outcomes serving as a critical benchmark for assessing prognosis and therapeutic efficacy following acute myocardial infarction (AMI) [1]. Apart from early reperfusion, there are no therapies available to reduce the extent of injury or promote cardiac regeneration. To address this unmet need, cardioprotective therapies are continuously investigated. However, despite numerous successful preclinical studies, translation of cardioprotective interventions remains elusive [2]. This discrepancy is attributed to several factors including the quality, analyzed comorbidities and comedications rigor, and transparency of preclinical studies [3], [4]. To overcome these issues, well-documented, rigorous multicenter studies with standardized protocols and analysis tools are needed [5], [6].

Quantifying the area at risk (AAR) and infarct size (IS) in preclinical models of myocardial infarction is essential for evaluating the effectiveness of cardioprotective interventions. Histological staining of heart slices with Evans Blue (EB) or similar stains to identify the AAR and with 2,3,5-triphenyltetrazolium chloride (TTC) to negatively stain the infarcted tissue, provide a robust, widely accepted method for assessing IS [7], [8].

Using image analysis software, such as ImageJ [9], researchers manually trace the myocardium, AAR, and IS to calculate key metrics such as the percentage of IS within the AAR and AAR within the myocardium. While this method is widely used in small animal experiments [10], [11], it is time-intensive and prone to observer bias. Furthermore, it lacks objective standardization, highlighting the need for advanced automation tools to optimize the process and improve reproducibility.

To address some of these issues, we previously developed a software specifically designed for planimetric IS analysis (Infarctsize™, Pharmahungary Group, Szeged, Hungary) which allowed for manual analysis and its archiving [12]. While this tool improved reliable data management, it did not simplify the analysis process, which thus remained complex, time-consuming, and subjective.

Artificial intelligence (AI) is increasingly being used for automated image analysis in both clinical and basic science settings. AI systems have been successfully applied to the analysis of clinical cardiac magnetic resonance- and computer tomography data to facilitate decision-making [13], [14], [15]. Furthermore, deep learning-based segmentation has been used for IS prediction with contrast-enhanced magnetic resonance imaging in rodents [16]. In addition, a deep learning image segmentation model was introduced in a preliminary study for preclinical IS quantification. However, the lack of validation across images from different laboratories, the non-disclosure of trained AI models, and the absence of a user interface hinders the public use of the tool without strong bioinformatics support [17].

To overcome the limitations of currently available tools, within the framework of the EU-CARDIOPROTECTION and IMPACT COST Actions (CA16225, IG16225), we aimed to develop an AI-based software for the segmentation of Evans Blue and TTC-stained heart slices to calculate AAR and IS, which will provide a convenient platform for IS image analysis together with data management capabilities to reduce analysis time and possibly decrease inter-observer variability for small rodent studies.

Here we present Infarctsize-AI™ (available at: https://infarctsize.com), a software that (1) implements AI analysis on Evans Blue and TTC-stained rodent heart slices, (2) supplies a Confidence Score on AI analysis to highlight areas with poor annotation, (3) provides an online, easy-to-use platform for manual supervision and correction that (4) includes data management capabilities essential for high-quality documentation and reporting necessary for reproducible and transparent research. The software has been developed in accordance with the international guidelines for AI-based cardiovascular analysis tools [18]. Here we demonstrate that the AI effectively analyze relevant areas on TTC and Evans Blue-stained heart slices, significantly reducing analysis time and inter-observer variability, without risking imperfect automated analysis.

## Materials and methods

### Image source and animal models

For the training of the neural networks, images from pre-existing research studies [4], [19], [20], [21], [22] were retrospectively analyzed. All of these earlier experiments adhered to the Directive 2010/63/EU of the European Parliament on the protection of animals used for scientific purposes and the Animal Research: Reporting of In Vivo Experiments (ARRIVE) guidelines [23]. The protocols were formally approved by the appropriate national or institutional ethics committees.

Animals were subjected to acute left anterior descending coronary artery ligation for 30 minutes in rats and 45 minutes in mice followed by reperfusion for 120 minutes or 24 hours in rats and 24 hours in mice. At the end of the experiments, hearts were excised and AAR was negatively stained with EB. Then the hearts were sliced and stained with TTC to visualize the infarcted area. Finally, images from both sides of the slices were taken [24].

### Training of deep learning segmentation models

Training dataset consisted of images of rat heart slices from two laboratories with 248 and 325 slice images and of images of mouse heart slices from one laboratory with 77 slice images. Training data had existing analysis for slice area, AAR and infarcted area. These analyses were used as reference during the training of the segmentation models. Images for the training were resized to 224 x 224 pixels. Horizontal and vertical flipping were employed during the training process as data augmentation.

Three neural networks were trained independently for the pixel-wise segmentation of the heart slice images. Two of these, U-Net++ [29] and DeepLabV3+ [30] were employed as pre-trained models from the segmentation_models.pytorch library [31]. In both cases, a ResNet34 backbone pretrained on the ImageNet dataset was used. The third neural network was a non-pretrained U-Net-based architecture. The encoder block of our U-net started with a double convolution, consisting of two 3×3 convolutional layers, each followed by batch normalization and a Leaky ReLu activation layer. This double convolution was repeated four times, always followed by a 2×2 max pooling layer. The decoder block consisted of four upsampling layers, each containing a transposed convolution operation and a double convolution. The final 3×3 convolution was repeated once more at the end of the decoder block.

Training was conducted using Adam optimizer [32], a learning rate of 0.006, a batch size of 16 and a loss function combining a sigmoid layer and binary cross-entropy loss. The training was stopped when the test loss did not significantly decrease for ten consecutive epochs. The training was executed in Google Colaboratory [33], using GPU servers provided by Google (Mountain View, CA, USA).

To generate a single combined output for the three neural networks, the probability of being part of the slice, AAR or infarct region were averaged pixelwise, respectively. Then pixels with an average probability greater than 0.5 were classified as part of the respective region. Combined output was used as the result of the AI analysis.

### Evaluation of AI analysis

One of the training sources provided 496 analyzed images of rat cardiac slices from 41 rats, which were not used for the training, as internal test data. Additionally, 122 analyzed rat heart slice images from two other laboratories (n = 60 and 62), which sources were not used for training, were employed as external test data. IS/AAR was calculated for each slice from AI analysis and manual analysis. Additionally, Overlap percentage was calculated as Jaccard Index × 100 between AI analysis and manual analysis for slice area, AAR and infarcted area. Slices were excluded from IS/AAR comparison if any of the analysis types did not provide AAR analysis. Slices were excluded from Ovelap percentage comparison, if none of the analyses provided annotation for the analyzed region type.

To compare AI analysis with manual analysis on whole-heart level, the total weight of AAR and infarcted area was calculated for each hearts based on region sizes and slice weights and then infarction weight was divided by AAR weight.

### Calculation and analysis of Confidence Score

Confidence Score was calculated by collecting and averaging the pixel probabilities of those pixels that had a probability over 0.5 in at least one neural network output. Finally, the values were stratified between 0 and 1.

To evaluate the performance of the Confidence Score, regions with an overlap below the pre-defined threshold of 75% in the comparison between AI analysis and manual analysis were categorized as low-reliability annotations for both AAR and infarct areas. Receiver operator characteristics (ROC) analysis was performed to assess whether Confidence Score serves as a predictor of low-reliability annotations.

### Experiments analyzing inter-observer variability and analysis time

Rat heart slice images from internal (n = 36) and external (n = 37) sources were analyzed by two evaluators either with AI assisted analysis, consisting of AI analysis followed by manual correction or with manual analysis. Analysis time for each slice was recorded for both analysis methods. Overlap percentage, defined as Jaccard Index × 100, was calculated between the evaluators in both analysis methods. Slices were excluded from Overlap percentage comparison, if none of the compared analyses provided annotation for the analyzed region type. IS/AAR was also calculated from all analyses for each slices. Slices were excluded from IS/AAR comparison if AAR was not provided by one of the compared analyses types (Figure 1.).

**Figure 1:**
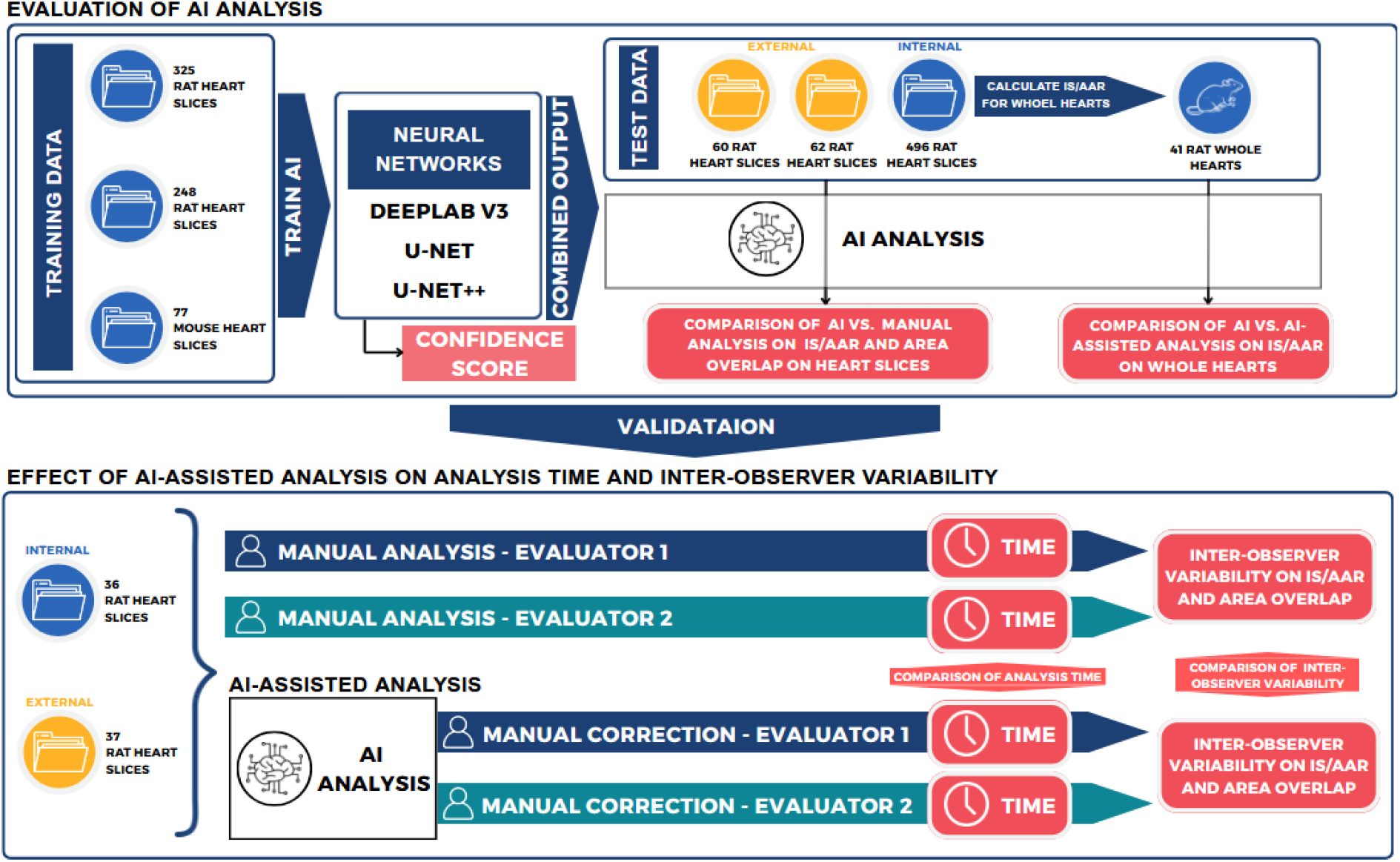
Study outline. Evans blue and TTC-stained Cardiac slice images from three independent laboratories were used to train neural networks with three architectures, the output of which were combined into a single AI analysis. Confidence score was calculated based on neural network outputs. AI analysis was tested on heart slices images from internal and external sources. Whole-heart data was also calculated and compared with AI analysis in the case of internal dataset. To validate AI analysis, two independent evaluators analyzed heart slice images from internal and external sources with or without AI analysis. Analysis time and inter-observer variability was investigated.

### Development of the user interface

The software consists of two components: a web browser-based user interface and a server that implements AI image analyis. The user interface is based on the VGG Image Annotator (VIA) version 2 [25], which has been extended using standard web technologies (HTML, CSS, and “vanilla” JavaScript programming language). To improve responsiveness and user interface aesthetics, the Bootstrap CSS framework [26] was integrated. Additionally, the Panzoom.js library [27] was employed for advanced interactive features, such as smooth zooming and panning, The server is implemented using Flask [28], a micro web framework written in Python, providing the backend functionality required for AI image analysis.

### Statistics and visualization

Statistical analysis was performed using R programming language [34]. Concordance Correlation Analysis was used to compare IS/AAR values between analysis methods. Lin’s Concordance Correlation Coefficient (CCC) and its 95% confidence interval (CI) are presented. The Wilcoxon signed-rank test was used to compare overlap between the two evaluators in the manual versus AI-assisted analysis, and to compare analysis time. Spearman’s rank correlation was used for the correlation analysis of Confidence Score and overlap on AAR and infarct regions. Figures were created using R programming language [34] and Canva application (canva.com).

## Results

### Infarctsize-AI™: web browser-based application for AI-assisted infarct size measurement

To facilitate the analysis of TTC- and EB-stained cardiac slices for IS measurement, we developed a web browser-based planimetric analysis software featuring AI-assisted analysis capabilities. The software prioritizes simplicity, ease of use, and rigorous reporting to support reproducible and efficient research workflows (Figure 2A).

**Figure 2:**
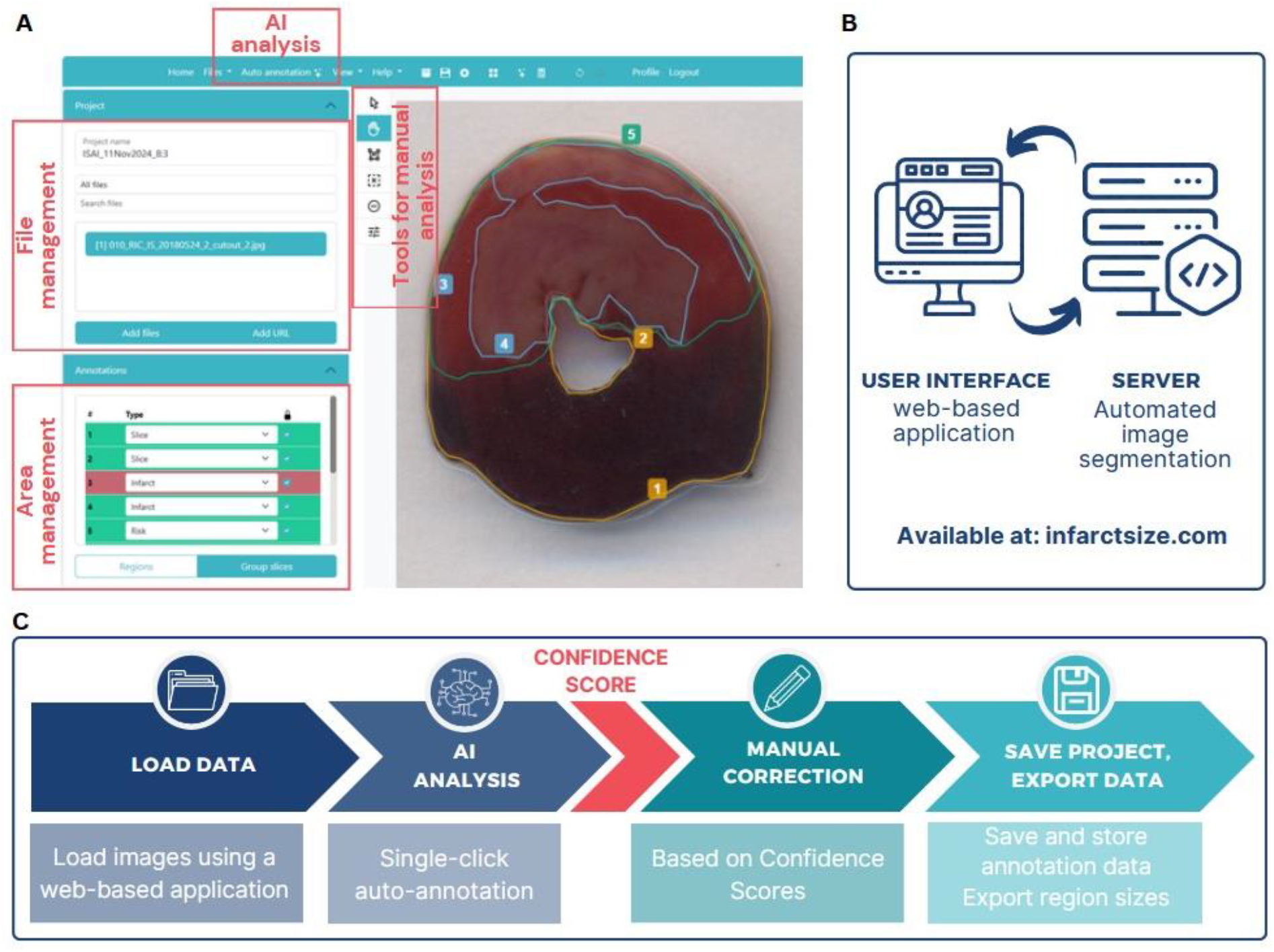
Infarctsize-AI is a web-based application for rodent infarct size measurement, that (A) implements image annotations with an ease-of-use toolkit and (B) enables automated image segmentation using AI analysis. (C) Images can be loaded into the application and after AI analysis, correction based on Confidence Score can be applied. Segmentation and numeric data are stored for documentation and revision.

A key innovation of the software is the Auto Annotation functionality, which performs the AI analysis of the loaded images using the trained AI algorithm (Figure 2B). For each area, Confidence Score is calculated and displayed with an adjustable threshold to highlight area with potentially low-reliability. Users can apply manual corrections when necessary, using an intuitive toolset. Results and annotations can be exported for quantitative analysis, documentation, publication, and future revision (Figure 2C).

### AI analysis highly overlaps with manual analysis

To assess whether deep learning-based automated image segmentation can be applied for the analysis of EB and TTC-stained cardiac slices, we compared IS/AAR values derived from AI or manual analysis. The AI was tested on cardiac slice images of cardiac slices originating from one of the training laboratories (internal data) as well as on images from independent sources (external data).

For the internal data, AI analysis demonstrated good correlation with manual analysis with a CCC of 0.844 with 95% CI of [0.814; 0.869] (Figure 3A, Figure S1). Absolute differences between the AI analysis and the manual analysis was −0.22% in average with a 95% limit of agreement of [-28.6%; 28.1%] (Figure 3B). However, the AI failed to predict IS/AAR on external data (Figure 3C, Figure S2). When we investigated overlap values, we identified that AI analysis was excellent for slice areas and that it gave good predictions for AAR on both internal and external datasets, however, it identified infarct areas acceptably only for the internal data, while failed to do so for the external data (Figure S3).

**Figure 3:**
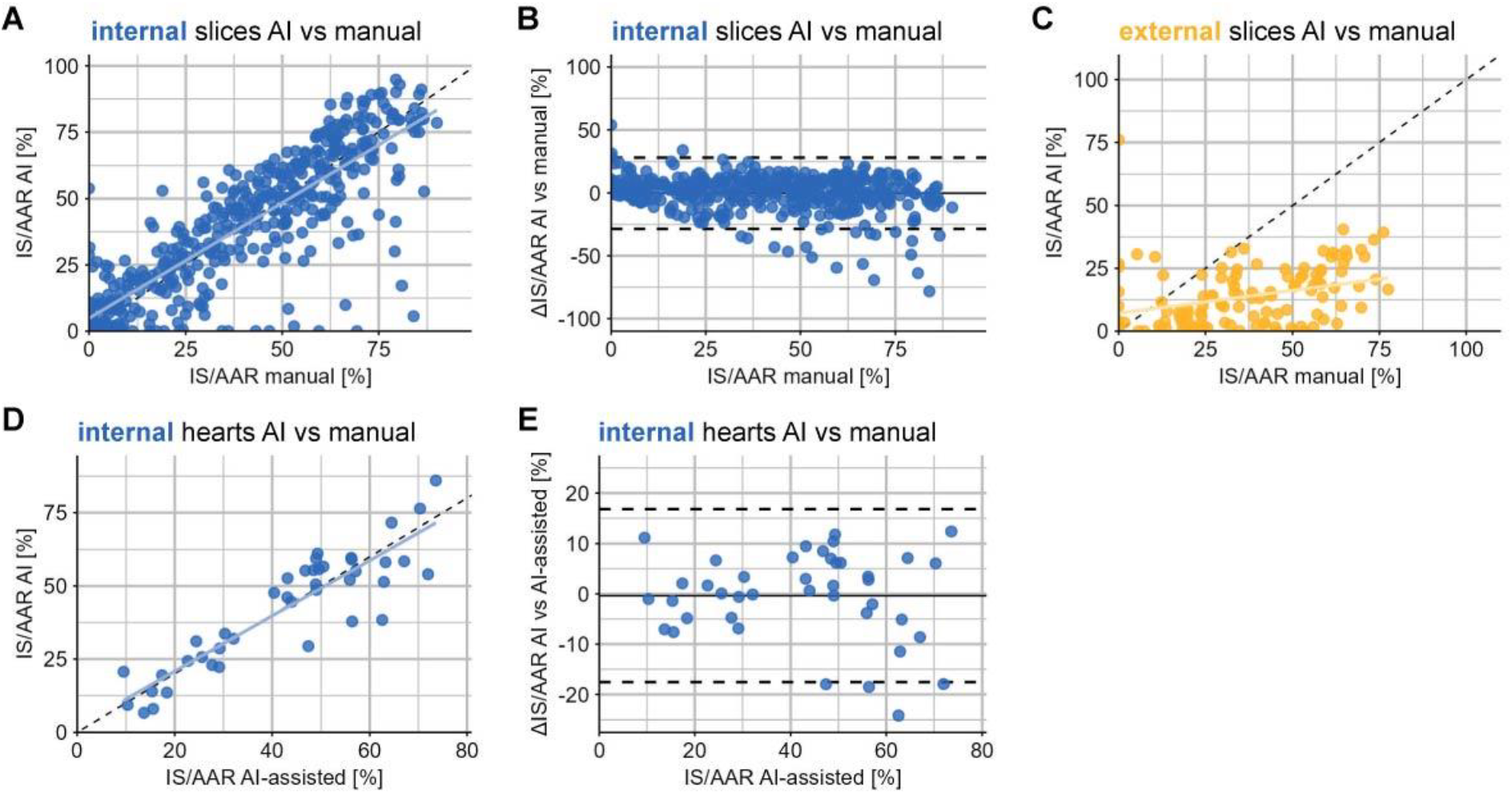
Investigation of the quality of AI analysis. (A) Correlation between AI analysis and manual analysis on internal dataset slices. (B) Differences between AI analysis and manual analysis on internal dataset slices. (n = 429) (C) Correlation between AI analysis and manual analysis on external dataset slices. (n = 120) (E) Correlation between AI analysis and manual analysis on whole-heart internal data (n = 41) (F) Differences between AI analysis and manual analysis on whole-heart internal data (n = 41).

We then compared AI analysis on internal data of whole hearts. Strong correlation was demonstrated by a CCC of 0.894 with 95% CI of [0.812; 0.942] (Figure 3D). Mean differences between AI analysis and AI-assisted analysis was −0.37% with the 95% limits of agreement of [-17.6%;16.8%] (Figure 3E).

These results indicate that AI analysis performs well in annotating slice areas and AAR on both internal and external data. In the case of internal data, AI analysis provides reliable annotations for infarct areas as well, resulting in minor errors in the final IS/AAR values. However, in some cases, AI analysis results in significant errors, therefore, human supervision over AI analysis is recommended.

### Confidence Score is a good predictor for AI analysis with low-reliability

To facilitate human supervision subsequent to the AI analysis, we aimed to develop a Confidence Score that is capable of identifying annotations with potentially low-reliability. We tested its ability to predict regions that have overlap below 75% between AI and manual analysis. As slice areas were annotated well in case of any tested data sources, we tested Confidence Score on AAR and Infarct areas only.

Using ROC analysis, we have identified that Confidence Score is an excellent predictor of overlap below 75% between manual and AI analysis, with balanced sensitivity and specificity. The area under the ROC curve was 0.968 for AAR (Figure 4A) and 0.954 for Infarct (Figure 4B) regions.

**Figure 4:**
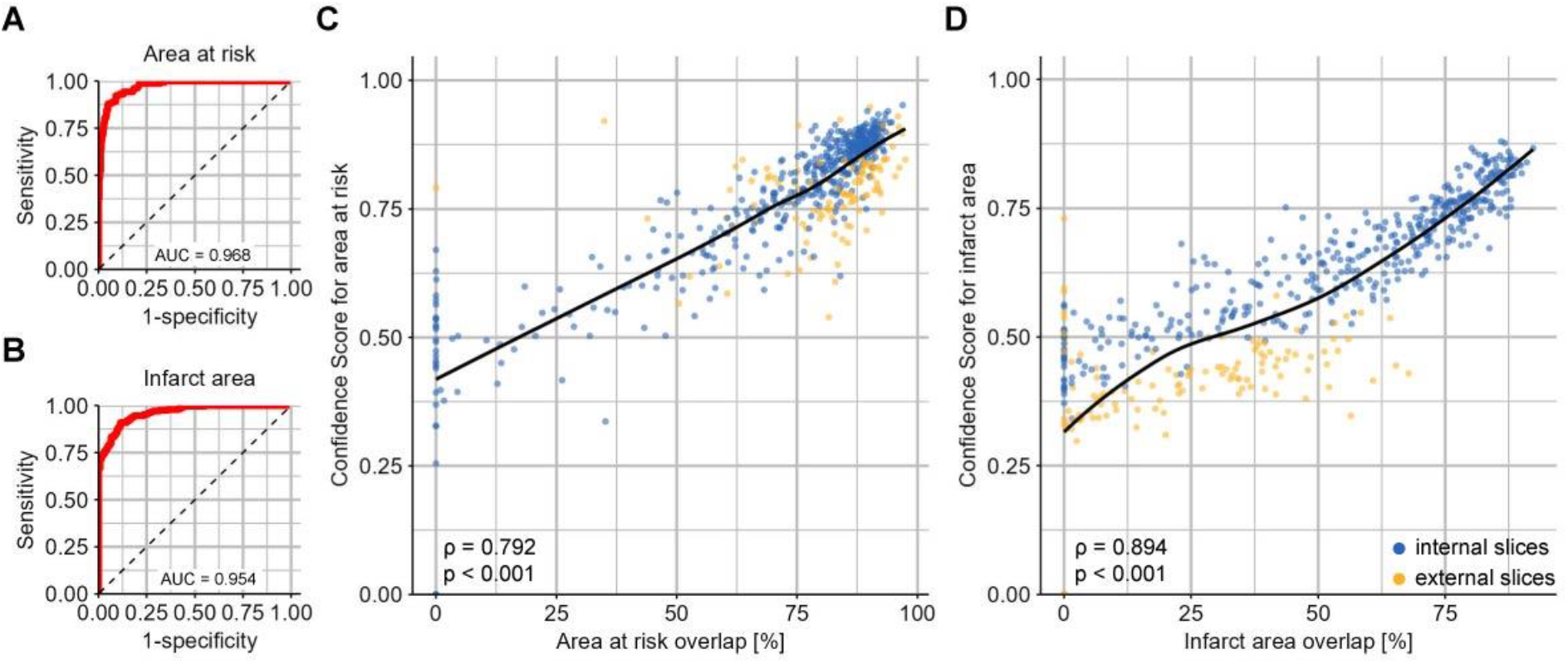
Analysis of Confidence Score for the identification of poor AI analysis defined as overlap < 75% (A,B) Receiver Operator Characteristics curves for (A) area at risk and (B) infarct regions. (C-D) Correlation analysis between overlap and Confidence Scores for (C) area at risk and (D) infarct regions.

We also conducted correlation analyses between the Confidence Score and region overlap. Results revealed strong correlation between Confidence Score and region overlap for both AAR (p < 0.001; ρ = 0.792) and infarct areas (p < 0.001; ρ = 0.894) (Figure 4C,D).

These results demonstrate that the developed Confidence Score discriminates between high- and low-reliability AI analysis effectively, making it suitable for highlighting annotations that require manual correction.

### AI-assisted analysis reduces analysis time and inter-observer variability

The primary goals of our AI-assisted analysis were to reduce analysis time and, if possible, inter-observer variability. To evaluate whether Infarctsize-AI™ can achieve these objectives, we conducted an experiment in which two independent evaluators analyzed the same images with or without AI analysis on internal and external heart slice images (Figure 1).

#### AI-assisted analysis reduces analysis time

We compared the time spent on the analysis of each slice by the evaluators with manual or AI-assisted analysis. We found that analysis time was greatly reduced on both internal and external datasets (Figure 5A,B). Average analysis time per slice was 271 seconds with manual and 139 seconds with AI-assisted analysis on internal dataset, which demonstrates a 49% reduction in analysis time.

**Figure 5:**
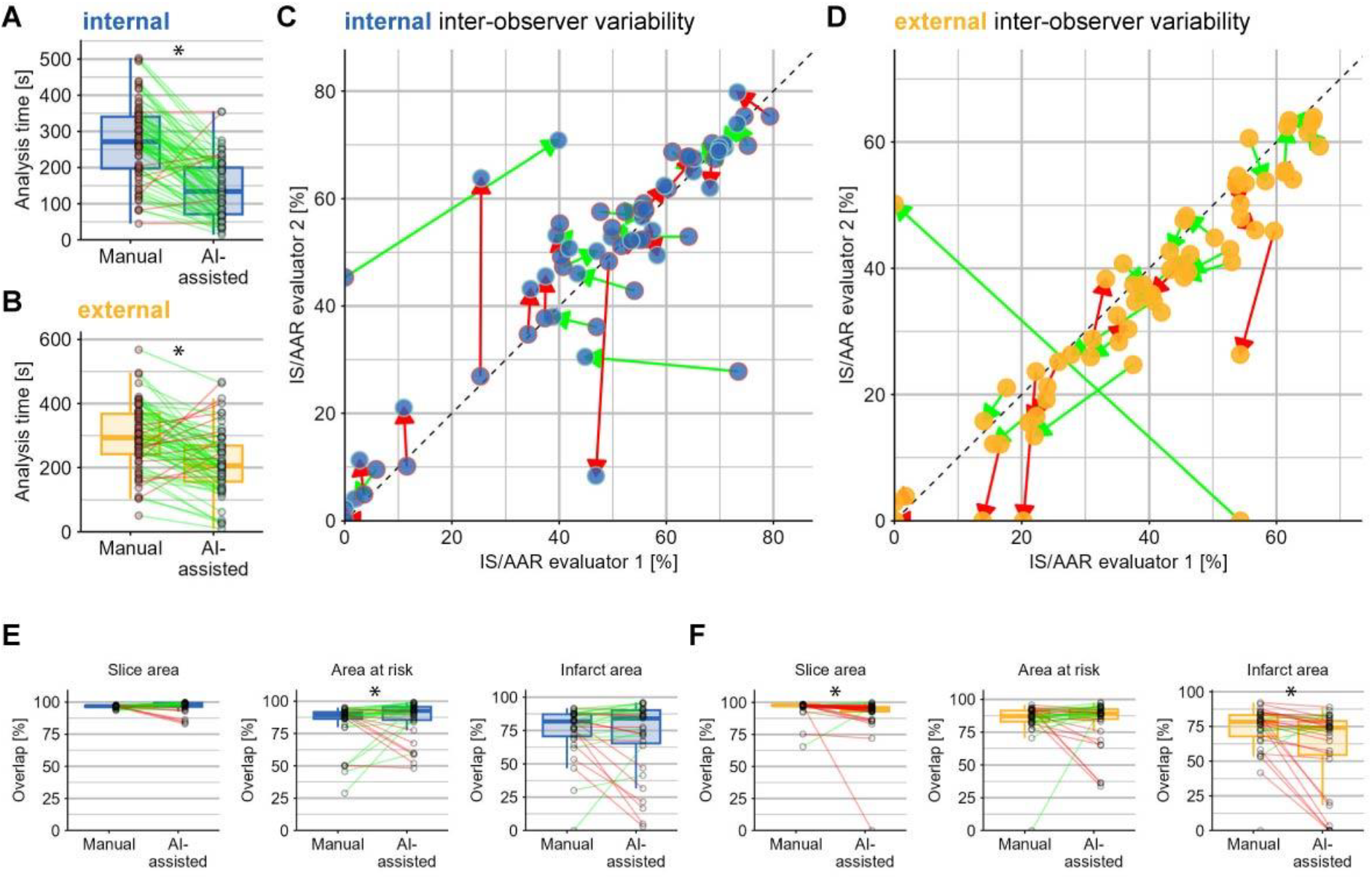
Assessment of the effect of AI-assisted analysis on analysis time and inter-observer variability (A-B) Time needed for image analysis with manual versus AI-assisted analysis on (A) internal (n = 73) and (B) external (n = 71) data (C-D) Correlation analysis between two independent evaluators on (C) internal (n = 35) and (D) external (n = 34) data. Data with manual or with AI-assisted analysis from the same heart slice are connected with arrows. Green arrows represent that difference between the two evaluators are reduced with the AI-assisted analysis. (E-F) Overlap values between the two evaluators with manual or AI-assisted analysis for slice, area at risk and infarct area for (E) dependent data and (F) independent data.^*^ p < 0.05 Wilcoxon signed-rank test.

#### AI-assisted analysis reduces inter-observer variability on internal dataset

To assess inter-observer variability, we compared CCC between the evaluators with or without AI-assisted analysis. We observed only minor increase with the AI-assisted analysis. (Internal data: manual CCC = 0.875 [0.769;0.935], AI-assisted CCC = 0.879 [0.774;0.937], External data: manual CCC = 0.838 [0.707;0.914], AI-assisted CCC = 0.847 [0.719;0.919])

When we compared overlap values between the two evaluators, we observed an increase by AI-assisted analysis for AAR in the case of internal dataset (Figure 5E). However, for external dataset, slice region and infarct region overlap was decreased (Figure 5F).

To further evaluate these results, we compared inter-observer variability with intra-observer variability. When we compared manual and AI-assisted analysis of the same evaluator, we observed that intra-observer CCC (Figure S4; intra-observer CCC: internal: 0.910 [0.858, 0.943], external: 0.849, 95% CI [0.769; 0.903]) was comparable with the inter-observer data. Similarly, within-evaluator overlap values were similar to those that were observed with manual inter-observer overlap. These results suggest that any approach that introduces manual correction could not further improve inter-observer variability, as it is proposed to be derived from the nature of the analysis other than disagreements between the evaluators.

## Discussion

In this report we introduce Infarctsize-AI™, an application designed for rodent IS measurement that enables AI analysis of heart slice images with possible manual correction. This tool provides a faster and more objective alternative than conventional manual analysis. Additionally, Infarctsize-AI™ offers capabilities for storing and exporting annotations of the specific areas for quality control and documentation.

AI analysis demonstrated strong agreement with manual analysis on whole-heart internal data, however, in certain cases major errors occasionally occurred. Therefore, we recommend manual supervision over AI analysis. As an assistance, we implemented a feature based on the Confidence Score which highlights regions with potentially low-reliability. As the Confidence Score has proven to be a good predictor for AI imprecision, we recommend limiting manual correction on the regions flagged by low Confidence Score. Such scores are recommended to be applied in various clinical settings as well, such as PET image analysis [35].

AI-based approaches demonstrated the possibility of reducing inter-observer variability in medical image analysis [36], [37]. Interestingly, we found only minor change in inter-observer variability, when we applied AI-assisted analysis. As our results highlighted that the degree of inter-observer and intra-observer variability are similar, we suggest that differences between evaluators are derived from the nature of the analysis and thus cannot be further decreased. If AI analysis was accepted without correction, inter-observer variability would be theoretically zero. However, AI differences are derived not only from the subjectivity of the analysis, but from errors as well, therefore, we recommend not to abandon manual supervision. Reports on evaluation of cancer MRI data by machine learning suggest that manual supervision over AI analysis is inevitable [38]. Since with the developed Confidence Score the identification of major AI analysis errors are possible, we recommend keeping AI analysis with high confidence scores unmodified and amending those areas only, where significant errors occurred. In this way, manual correction would bring in inter-observer variability only in those cases when the AI produced major error.

One limitation of our current AI tool is its performance on images from laboratories that were not included in the training. In these cases, AI analysis performed well for the analysis of the heart slice area and the AAR, but analysis on the infarcted area were inaccurate. Therefore, on these datasets, manual revision is necessary for all heart slices. However, our results indicate that AI-assisted analysis still reduces total analysis time. Multicenter studies demonstrate that even with high methodological agreement, major between-center differences occur in preclinical MI experiments [6], [24]. Therefore, a generalized AI model, which has equal accuracy on any dataset, may not be achievable.

For general accessibility of our software, we have developed a fully functional web-browser-based user interface, that integrates AI analysis, highlights poor AI analysis based on Confidence Score, provides a toolset for manual analysis or correction, and supports data management for documentation and revision. Using Infarctsize-AI™, the analysis of rodent Evans Blue and TTC-stained infarct size images becomes more objective, faster, and better documented. Therefore, our tool is well-suited for facilitating translatability of basic research in cardioprotection. Infarctsize-AI™ can be accessed at https://infarctsize.com.

## Author contributions

CK: conceptualization, methodology, formal analysis, investigation, writing-Original Draft, Project administration; DK: Formal analysis, Writing – Original Draft, Visualization; BYW: Investigation; TGG: Investigation; GBB: Investigation; BA: Methodology, Supervision; CT: Software; AR: Software, Formal analysis; AH: Conceptualization, Methodology, Resources, Supervision; SH: Investigation; DJH: Resources, Funding Acquisition; RV: Resources, Supervision; MO: Investigation; MW: Investigation; MM: Resources, Supervision; AM: Investigation; TS: Investigation; PB: Resources, Supervision, Funding Acquisition; TK: Resources; JI: Resources; RS: Resources; CJZ: Resources; IA: Resources; BKP: Resources; PF: Conceptualization, Resources, Writing - Review & Editing, Funding Acquisition; ZG: Conceptualization, Resources, Writing - Review & Editing, Funding Acquisition, Supervision; All authors revised the manuscript. All authors read and approved the final manuscript.

## Data availability

The software presented is available at: https://infarctsize.com.

Source codes are available at: https://dev.itk.ppke.hu/infarctsize-ai

All other data are available from the corresponding author upon reasonable request.

## Ethics approval

Animal data were collected from earlier, pre-exisiting studies, which were approved by the responsible national offices of the authors, including permissions of the National Scientific Ethical Committee on Animal Experimentation and the Semmelweis University’s Institutional Animal Care and Use Committee (H-1089 Budapest, Hungary) and the government of Food Chain Safety and Animal Health Directorate of the Government Office for Pest County (project identification code: PE/EA/1912-7/2017; date of approval: November 2017), Permissions Nr 36 and Nr 84, issued by Food and Veterinary Service Republic of Latvia, and the SingHealth IACUC (#2020/SHS/1563) and the Animal Research: Reporting of In Vivo Experiments (ARRIVE) guidelines.

