## supplementary figures for "Infarctsize-AI: an efficient infarct size image analysis tool for small rodent myocardial infarction studies"

Supplementary material

Csenger Kovácsházi[1,2], Dóra Kapui[1,2], Bennet Y Weber[1,2], Tamás G Gergely[1,2], Gábor B Brenner[1,2], Bence Ágg[1,2,3], Csanád Tabajdi[4], Adrienn Rácz[4], András Horváth[4], Sauri Hernandez-Resendiz[5,6], Derek J Hausenloy[5,6,7,8], Reinis Vilskersts[9,10], Marta Oknińska[11], Michał Waszkiewicz[11], Michal Mączewski[11], Arnold Molnár[3], Tamara Szabados[12], Péter Bencsik[12,3], Thomas Krieg[13], Javier Inserte[14], Rainer Schulz[15], Coert J Zuurbier[16], Ioanna Andreadou[17,†], Bruno K Podesser[18], Péter Ferdinandy[1,2,3,*,#], Zoltán Giricz[1,2,3,*,#]

[1] Department of Pharmacology and Pharmacotherapy, Semmelweis University, Budapest, Hungary.
[2] Center for Pharmacology and Drug Research & Development, Semmelweis University, Budapest, Hungary.
[3] Pharmahungary Group, Szeged, Hungary.
[4] Faculty of Information Technology and Bionics, Pázmány Péter Catholic University, Budapest, Hungary
[5] Cardiovascular and Metabolic Disorders Programme, Duke-NUS Medical School, Singapore, Singapore.
[6] National Heart Research Institute Singapore, National Heart Centre Singapore, Singapore, Singapore
[7] Yong Loo Lin School of Medicine, National University Singapore, Singapore, Singapore
[8] The Hatter Cardiovascular Institute, University College London, London, UK
[9] Latvian Institute of Organic Synthesis, Riga, Latvia
[10] Faculty of Pharmacy, Riga Stradiņš University, Riga, Latvia
[11] Department of Clinical Physiology, Centre of Postgraduate Medical Education, Warsaw, Poland
[12] Cardiovascular Research Group, Department of Pharmacology and Pharmacotherapy, Albert Szent-Györgyi Medical School, University of Szeged, Szeged, Hungary.
[13] Department of Medicine, University of Cambridge, Cambridge, United Kingdom.
[14] Department of Cardiology, Vall d'Hebron University Hospital and Research Institute, Universitat Autònoma, Barcelona, Spain.
[15] Institute of Physiology, Justus Liebig University Giessen, Giessen, Germany
[16] Laboratory of Experimental Intensive Care and Anesthesiology (L.E.I.C.A.), Department of Anesthesiology, Amsterdam Cardiovascular Sciences, Amsterdam UMC, University of Amsterdam, The Netherlands
[17] National and Kapodistrian University of Athens, Faculty of Pharmacy, Department of Pharmaceutical Chemistry, Laboratory of Pharmacology, Panepistimiopolis, Zografou, Athens, Greece
[18] Ludwig Boltzmann Institute for Cardiovascular Research, Center for Biomedical Research, Medical University of Vienna, Vienna, Austria
[†] Deceased
* These authors are considered joint senior authors

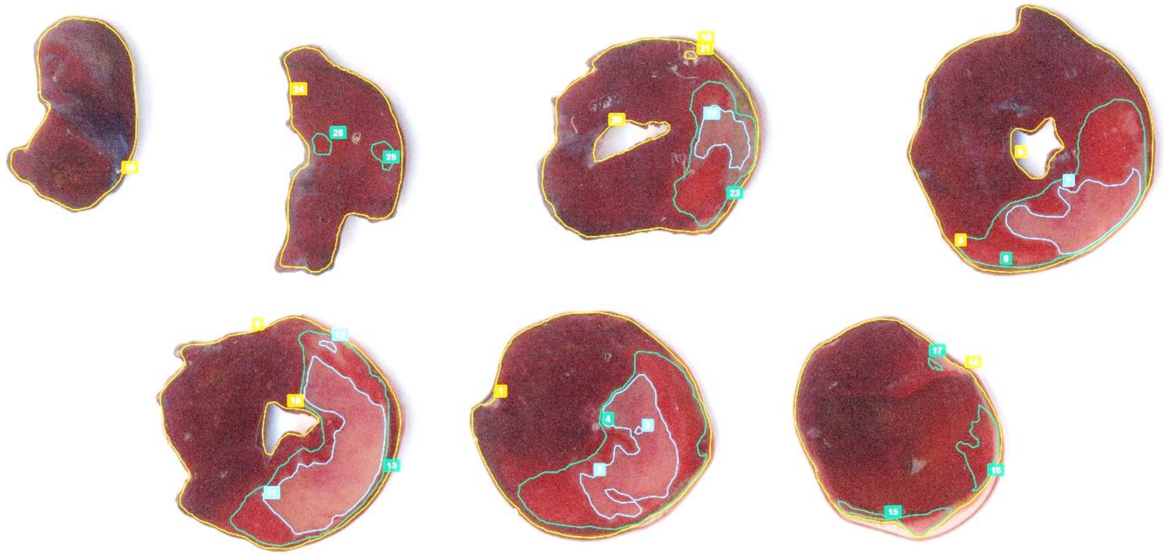

***Figure S1.:*** *Representative image of AI analysis on an internal data slice set image. Orange: slice area, Green: area at risk, Blue: infarct area*

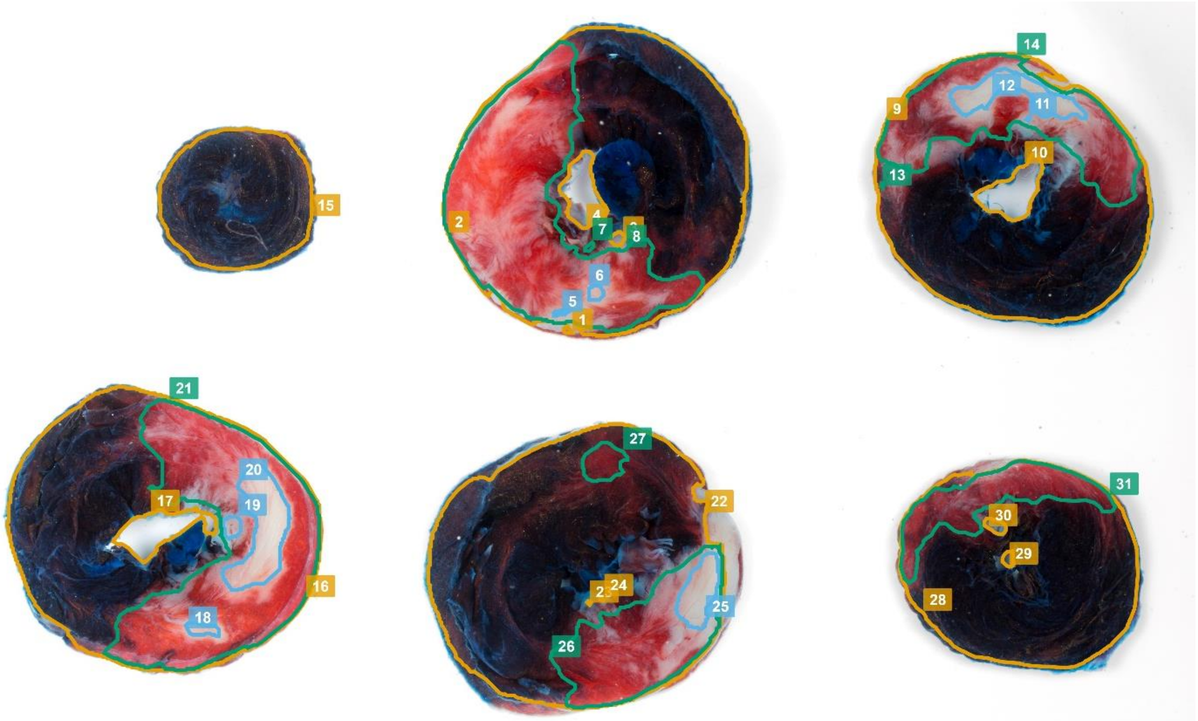


***Figure S2.:*** *Representative image of AI analysis on an external data slice set image. Orange: slice area, Green: area at risk, Blue: infarct area*

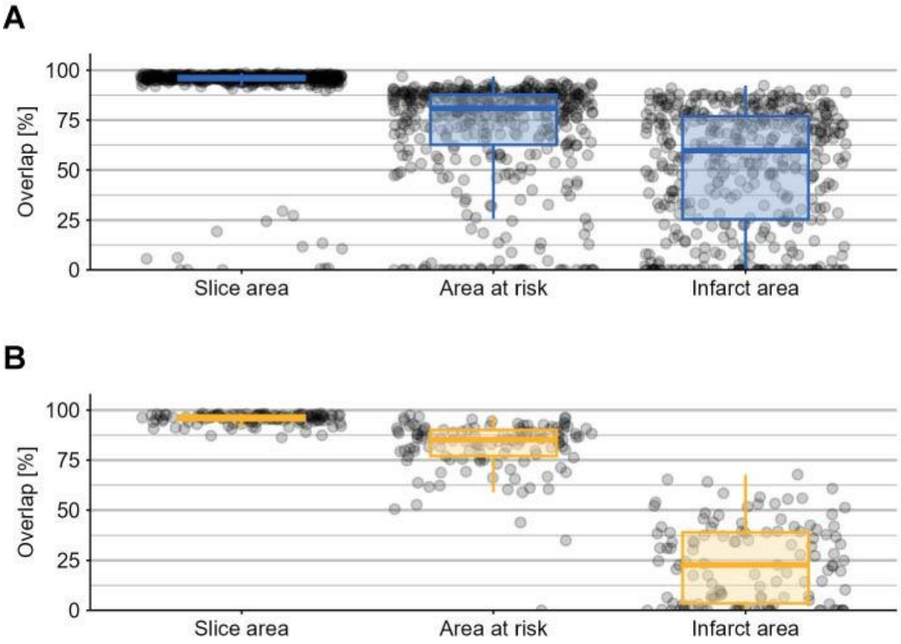


***Figure S3.*** *Overlap values for slice area, area at risk and infarct area between AI annotation and manual analysis on (A) internal and (B) external datasets*

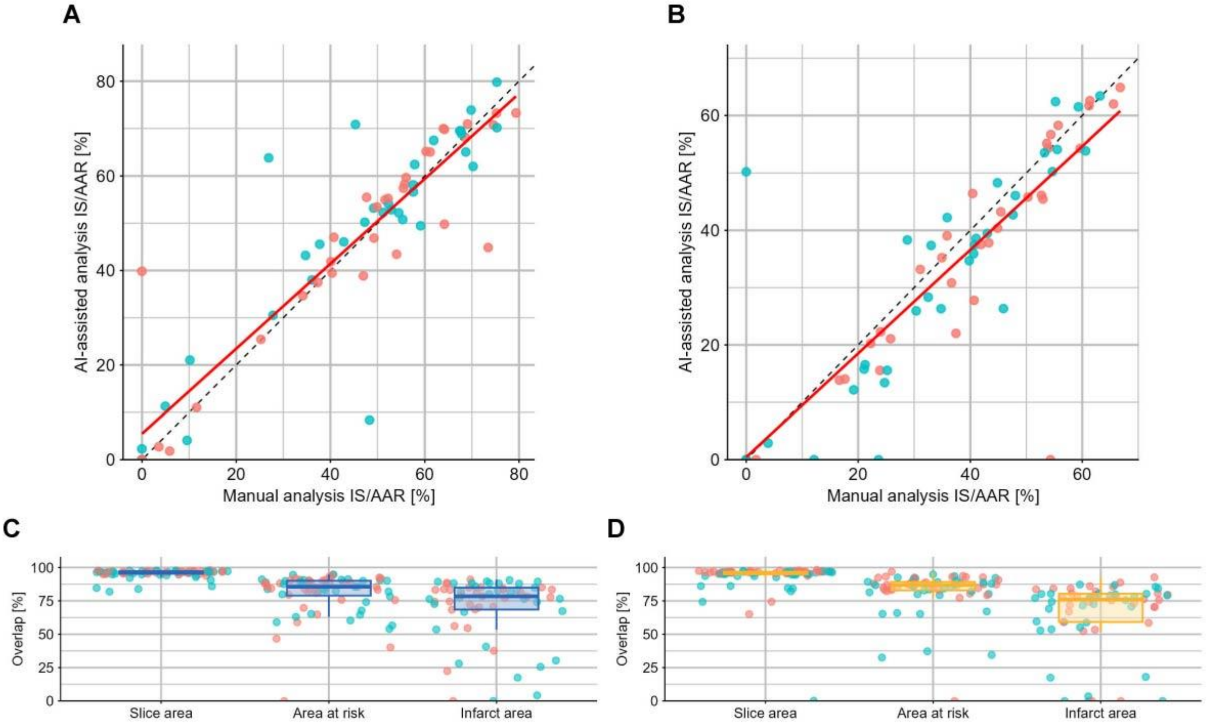


***Figure S4.****: Intra-observer variability. (A-B) Correlation between manual analysis and AI-assisted analysis within evaluator on (A) internal data and (B) external data. (C-D) Overlap between manual and AI-assisted analysis within evaluator on (C) internal data and (D) external data. Colors represent the two independent evaluators.*
